# Coalescent-based phylogenetic inference from genes with unequivocal historical signal suggests a polytomy at the root of the placental mammal tree of life

**DOI:** 10.1101/423996

**Authors:** Filipe R. R. Moreira, Carlos G. Schrago

**Affiliations:** Department of Genetics, Federal University of Rio de Janeiro, Rio de Janeiro, Brazil

**Keywords:** Phylogenomics, Placental root, Polytomy, Phylogenetic information

## Abstract

Finding the correct place of the root of the Placentalia tree of life is an unsolved problem in mammalian systematics. Three major competing hypotheses were proposed, alternating the position of the supraordinal taxa Afrotheria, Xenarthra, or Atlantogenata as sister lineages to the remaining placentals. While all three phylogenetic associations were inferred by studies conducted so far, recent assessments applying heterogeneous models and coalescent-based phylogenetic methods found robust support for the Atlantogenata hypothesis. Current developments in theoretical phylogenetics, such as the demonstration that concatenating genes may bias the likelihood function, and that summary coalescent-based phylogenetic methods are sensitive to errors in gene tree estimation, calls for a reevaluation of the early placental split problem. We investigated the phylogenetic relationships between placental superorders by restricting the analysis to subsets of genes with unequivocal phylogenetic signal. In contrast to previous works, we show that the hypothesis of a polytomy at the placental root could not be rejected under the multispecies coalescent model. This result endorses conclusions from analyses of retroposon insertion patterns. We provide an analytical framework to access empirical polytomies employing summary coalescent methods and topological tests, helping the investigation of poorly resolved nodes in the tree of life.

## Introduction

Rooting the placental tree of life is a major challenge in mammalian systematics (Teeling and Hedges, 2013). Three hypotheses are generally considered regarding the phylogenetic association of the major mammalian lineages, positioning Afrotheria (e.g. elephants and hyraxes; (Asher, 2007; Mccormack et al., 2012; Murphy et al., 2001; Nikolaev et al., 2007; Nishihara et al., 2007; Romiguier et al., 2013), Xenarthra (e.g., armadillos and sloths; (Kriegs et al., 2006; O’Leary et al., 2013; Waddell et al., 2001), or Atlantogenata, i.e., Afrotheria + Xenarthra (Hallström et al., 2007; Hallström and Janke, 2008; Kjer and Honeycutt, 2007; Meredith et al., 2011; Morgan et al., 2013; Murphy et al., 2007; Prasad et al., 2008; Song et al., 2012; Tarver et al., 2016; Wildman et al., 2007), as the sister group to the remaining placentals (**Figure 1**). So far, studies employing different data sources and methods, including genomic data, were unable to unambiguously resolve the early placental split, motivating the proposition that the first divergences of the placental tree were not strictly bifurcating (Churakov et al., 2009; Hallström and Janke, 2010; Nishihara et al., 2009). Alternatively, other works suggested that this phylogenetic incongruence may be attributed to the effects of incomplete lineage sorting (ILS) (Song et al., 2012), long branch attraction (LBA) (Romiguier et al., 2013), and model misspecification (Morgan et al., 2013; Tarver et al., 2016). Notwithstanding, assessments of this topological conflict adopting heterogeneous substitution models (Morgan et al., 2013; Tarver et al., 2016) and summary coalescent methods (Song et al., 2012; Tarver et al., 2016) found robust support for the Atlantogenata hypothesis, which was also recovered when previously published datasets were re-examined with best-fitting models (Tarver et al., 2016).

**Figure 1.**
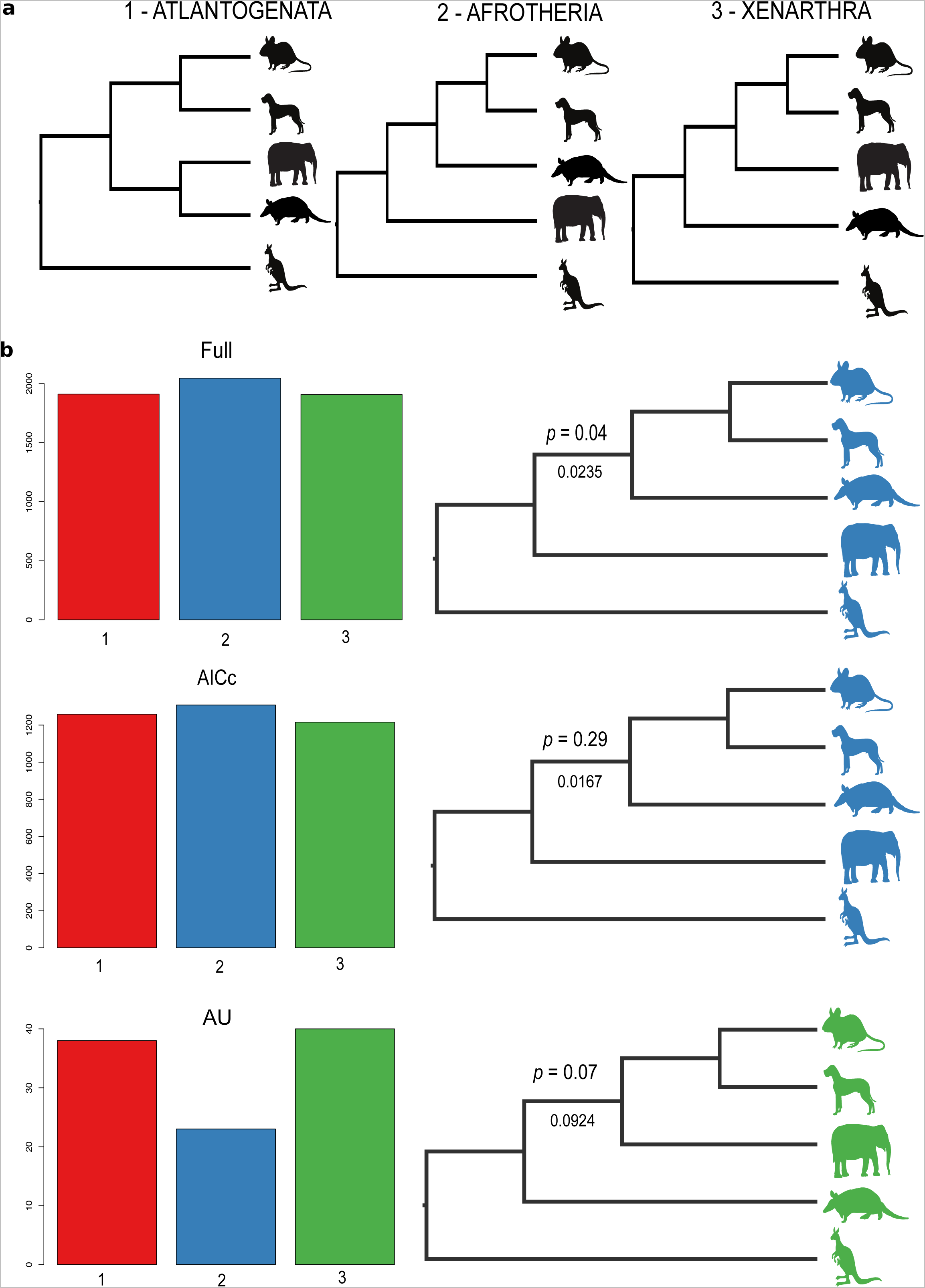
The three phylogenetic hypotheses for the position of the root of placental mammals (a). Distribution of gene tree topologies in the *full*, *AICc*, and *AU* datasets, with their respective species tree resolution inferred with Astral-III. The *p*-value of the polytomy test (above) and lengths in coalescent units (bellow) were given for branches responsible for resolving the polytomy at the root (b).

Despite these results, controversy remains, as methods employed in previous analyses are not free of biases from different sources. For instance, it has been shown that, when sequences are concatenated for carrying out supermatrix analysis, only a small fraction of sites may be responsible for choosing between alternative topologies (Shen et al., 2017). Moreover, it has been demonstrated that the likelihood function used in phylogenetics is inappropriate when genes are concatenated, resulting in positively misleading estimates under high levels of gene trees/species tree incongruence (Degnan and Rosenberg, 2006; Kubatko and Degnan, 2007; Mendes and Hahn, 2018; Roch and Steel, 2015), which might be the case of the early placental split (Song et al., 2012; Tarver et al., 2016). On the other hand, methods that avoid concatenation by accounting for the long-term effects of ILS were criticized in the context of placental phylogeny (Springer and Gatesy, 2016). The criticism lies on the fact that summary coalescent-based phylogenetic methods treat gene trees as error-free observations (Song et al., 2012). As maximum likelihood trees are point estimates, treating them as observations imply overconfidence, which may bias phylogeny estimation by approaches derived from the multispecies coalescent (MSC) model (Xi et al., 2015). This issue has been largely ignored by analyses of the placental tree relying on coalescent-based methods (Mccormack et al., 2012; Song et al., 2012; Tarver et al., 2016). It is thus conceivable that previous studies on placental phylogeny were impacted by genes with ambiguous signal for resolving the early divergences.

Following this reasoning, in this work, we were prompted to investigate, under the MSC model, which phylogenetic hypothesis regarding the early placental split was significantly favored when only subsets of phylogenetically informative genes were employed. We examined the mammalian phylogeny from two subsets of genes trees that either presented unequivocal phylogenetic resolution for the early placental split or consisted of topologies that did not overfit data.

## Material and Methods

Our analysis was carried out using 14,526 orthologous coding sequences retrieved from the OrthoMam v9 data base (Douzery et al., 2014), which were filtered to consist solely of genes whose maximum likelihood (ML) tree contained an outgroup and at least one representative of each placental superorder, namely, Xenarthra, Afrotheria, Laurasiatheria and Euarchontoglires. Furthermore, to reduce the phylogenetic problem to a 4-terminals tree, which makes the topological space computationally accessible, only ML trees wherein superorders were recovered as monophyletic were kept. We thus focused on the phylogenetic resolution of the early placental divergence, allowing us to evaluate all the three possible topological hypotheses (**Figure 1**). After this filtering step, our dataset was composed of 5,862 alignments, hereafter referred to as the *full* dataset.

Gene tree estimation and choice of substitution models were performed with the IQ-TREE software (Kalyaanamoorthy et al., 2017; Nguyen et al., 2014). The statistical support for the three topological configurations at the root was assessed by the approximately-unbiased (AU) test (Shimoidara, 2002), also implemented in IQ-TREE. Genes with unequivocal phylogenetic signal, i.e., those with exclusive support for a single topology, were grouped, constituting the *AU* dataset. Finally, we also built a dataset encompassing genes that, under the corrected Akaike information criterion (AICc) (Hurvich et al., 1998), rejected the hypothesis of a polytomy between the mammalian superorders, suggesting a strictly bifurcating tree as the best fit model. Species tree inference was conducted independently for each dataset (*full*, *AU*, and *AICc*) employing the coalescent-based algorithm implemented in the Astral-III software (Zhang et al., 2017), which also performs hypothesis testing of polytomies according to the MSC model expectation (Sayyari and Mirarab, 2018). A significance level of 5% was used throughout this study. Astral was also used to estimate branch lengths in coalescent units, which is relevant for interpreting gene tree distribution in light of the MSC, in particular to investigate whether the branching pattern that defines the deep splits between the major lineages of placentals lies in the anomaly zone (Degnan and Rosenberg, 2006).

## Results

The distribution of ML gene trees supporting each of the three possible phylogenetic hypotheses of the early placental split was approximately equivalent in the *full* dataset. Out of 5,862 gene trees, 1,910 (32.6%) supported the Atlantogenata hypothesis, whereas 2,045 (34.9%) supported Afrotheria, and 1,907 (32.5%) favored the Xenarthra tree. Therefore, the majority of gene trees, as well as MSC-topology from Astral analysis, supported the Afrotheria tree. The coalescent-based polytomy test also rejected the null hypothesis of a polytomic topology for this distribution of gene trees (*p* = 0.04). However, when each gene from the *full* dataset was subjected to the AU test, we found that only 101 genes (0.02%) presented unequivocal phylogenetic signal for a single topology concerning the early placental split (the *AU* dataset). Each of the three possible topological associations were represented with the following frequencies among these gene trees: 37.6% supporting Atlantogenata, 22.8% Afrotheria and 39.6% Xenarthra. When Astral analysis was run using the *AU* dataset, the Xenarthra topology was inferred as the species tree, although the null hypothesis of a polytomy at the root of placental mammals was not rejected (*p* = 0.08).

According to the AICc test, 2,079 gene trees, nearly 35% of the *full* data set, overfit data. The distribution of topologies inferred from the remaining 3,782 genes that favored a strictly bifurcating tree as the best fit model (the *AICc* dataset) presented a similar frequency of each early placental phylogenetic hypothesis, namely, Atlantogenata: 1,259 (33.3%), Afrotheria: 1,308 (34.6%), and Xenarthra: 1.216 (32.2%). Astral inferred the Afrotheria as the placental species tree using the *AICc* dataset, but the hypothesis of a polytomic association among superorders was also not rejected (*p* = 0.29). Branch lengths establishing the pattern of the early placental split in the *full*, *AU*, and *AICc* datasets measured 0.0235, 0.0924, and 0.0167 coalescent units respectively (**Table 1**).

**Table 1.**
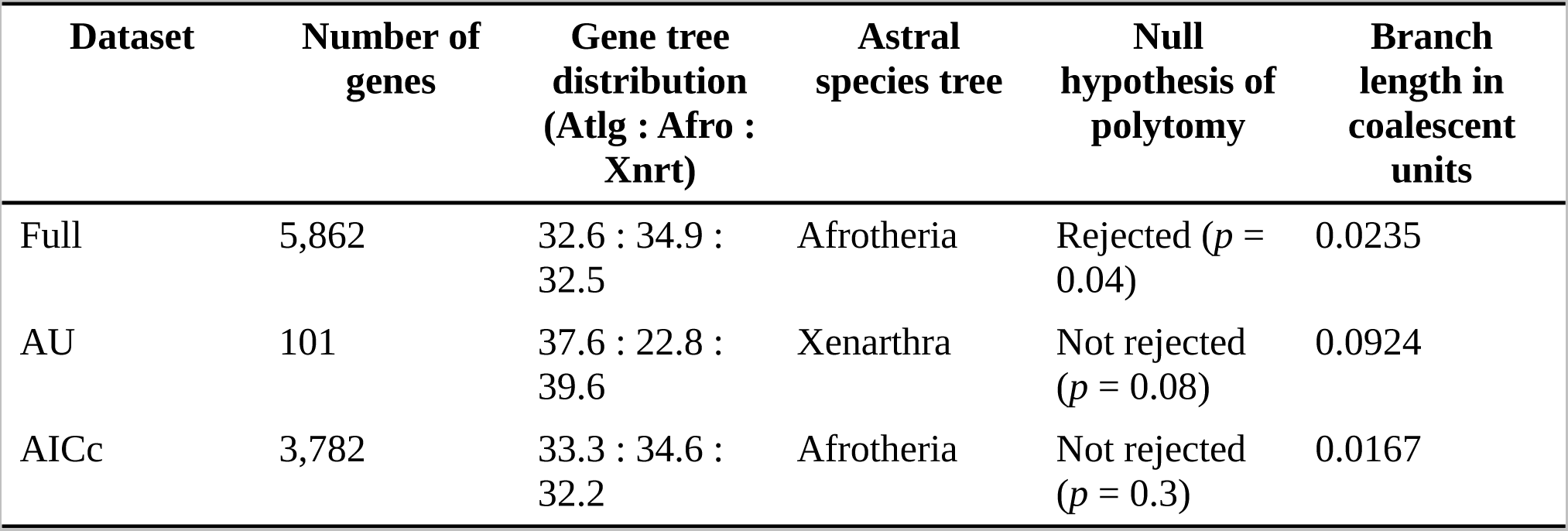
Summary of results from analyses employing three different filters on the alignments from OrthoMam.

## Discussion

Our results indicated that when genes with significant phylogenetic signal were used, Afrotheria or Xenarthra were the likely sister lineages of the remaining placentals. However, in both datasets, the hypothesis of a polytomy at the root of the placental tree was not rejected. When it comes to phylogenomics, all three phylogenetic hypotheses depicted in **Figure 1** were obtained by combining different methods and datasets. For instance, support for the Afrotheria tree was found when datasets less prone to LBA artifacts were employed (Romiguier et al., 2013), and when ultraconserved elements were analyzed (Mccormack et al., 2012). With a single exception (Kriegs et al., 2006), the Xenarthra tree has been rarely recovered by molecular data, although it stands as the preferred hypothesis of morphologists (O’Leary et al., 2013). In our analyses, we found no support for the Atlantogenata tree. This hypothesis was recently endorsed by Tarver *et al.* 2016, who implemented MSC methods and argued that correcting for heterogeneous substitution processes supports Atlantogenata as sister lineage to the remaining placentals. On the other hand, most previous works have relied on concatenated supermatrices (Hallström et al., 2007; Hallström and Janke, 2008; Mccormack et al., 2012; Morgan et al., 2013; Nikolaev et al., 2007; Nishihara et al., 2007; Romiguier et al., 2013; Wildman et al., 2007), which may be affected by the few-sites/large-effect issue (Shen et al., 2017). In face of these conflicting results, our study indicates that this lack of resolution for the early placental split might be due to the failure of the distribution of topologies from genes with significant phylogenetic signal to reject a polytomic association among mammalian superorders.

In this sense, it is meaningful to investigate whether the branching pattern of the early placental split lies in the anomaly zone, the region of parameter space where the most common gene tree is incongruent with the species tree (Degnan and Rosenberg, 2006). Even though it has been argued that the anomaly zone is unlikely to be a problem when studying empirical datasets (Huang and Knowles, 2009), the branch lengths supporting resolutions for the early placental split in all species trees inferred here were shorter than ~0.156 coalescent units (**Table 1**). Therefore, the theoretical possibility of obtaining anomalous gene trees distribution exists when investigating supraordinal relationship of placental mammals. Under this scenario, phylogenetic inference from concatenated sequences will be biased (Degnan and Rosenberg, 2006; Kubatko and Degnan, 2007; Mendes and Hahn, 2018; Roch and Steel, 2015).

By restricting our analysis to subsets of genes with significant phylogenetic signal, our study also avoided setbacks of summary coalescent methods (e.g., Astral) when gene tree uncertainty is not accounted for. It is worth noting that, although Tarver *et al.* (2016) implied that few genes have unequivocal signal for resolving the placental root by likelihood comparisons, which we corroborated via proper topological tests, the authors did not restrict their MSC phylogeny inference to this subset of unequivocal genes. Instead, they controlled for errors in gene trees by filtering topologies with non-parametric bootstrap thresholds (Felsenstein, 1985). The bootstrap, however, is not a formal statistical test between alternative topologies, and such procedure may have resulted in topological overconfidence.

The hypothesis of a polytomy at the root of placentals has been previously advocated by studies relying on retroposon insertions as phylogenetic markers (Churakov et al., 2009; Nishihara et al., 2009). Alternatively, using supermatrices, Hallström and Janke (2010) argued that the phylogenetic relationship among placental superorders is best represented by a phylogenetic network. While the proposition of a placental root polytomy is not entirely novel, this is the first coalescent-based analysis supporting this view, contradicting previous phylogenomic studies that were possibly affected by the very issues that motivated our reexamination of this problem (Hallström et al., 2007; Hallström and Janke, 2008; Mccormack et al., 2012; Morgan et al., 2013; Nikolaev et al., 2007; Nishihara et al., 2007; Romiguier et al., 2013; Song et al., 2012; Tarver et al., 2016; Wildman et al., 2007). Our results were not able to devise whether the placental root is the case of a soft (dataset limited) or hard (real) polytomy (Whitfield and Lockhart, 2007), but we showed how the prior examination of the historical signal of genes, associated with a control for systematic biases, impacts the resolution of a hard phylogenetic problem.

As conservative as our approach was, it illustrated how the MSC model may be used to test for polytomies in empirical datasets (Edwards, 2009). Future application of this framework is promising in unveiling rapid radiations in the deeper nodes of the tree of life (Whitfield and Lockhart, 2007). As has been argued by Shen et al. (Shen et al., 2017), branches that fail to reject a polytomy should be considered unresolved, implying a more accurate view of the signal available in data. We here provided evidence that this is the case of the early split between superorders of Placentalia.

## Acknowledgements

This research was supported by grants 310974/2015-1 and 440954/2016-9 from the Brazilian Research Council (CNPq) to CGS.

